# Rapid Motor Adaptation via Population-level Modulation of Cerebellar Error Signals

**DOI:** 10.1101/2024.01.03.574031

**Authors:** Vinh Nguyen, Brandon M. Stell

## Abstract

A core principle of cerebellar learning theories is that climbing fibers from the inferior olive convey error signals about movement execution to Purkinje cells in the cerebellar cortex. These inputs trigger synaptic changes which are purported to drive progressive adjustment of future movements. Individually, binary complex spike signals lack information about the sign and magnitude of errors which presents a problem for cerebellar learning paradigms exhibiting fast adaptation. Using a newly developed behavioral paradigm in mice, we introduced sensorimotor perturbations into a simple joystick pulling behavior and found parasagittal bands of Purkinje cells with reciprocal modulation of complex spike activity. Whereas complex spiking showed little modulation in the unperturbed condition, alternating bands were activated or inhibited when the perturbation was introduced, and this modulation encoded the sign and magnitude of the resulting sensorimotor mismatch. These findings provide an important piece of information for the understanding of cerebellar learning that helps to explain how the cerebellum could use supervised learning to quickly adapt motor behavior in response to perturbations.

**One-Sentence Summary:** Populations of Purkinje cells in the cerebellum facilitate trial-by-trial movement refinement by converting binary signals from movement errors into graded signals that encode the error’s direction and magnitude.

## Main Text

Established theories of cerebellar learning propose that climbing fibers (CF) signal performance errors in Purkinje cells, leading to depression of parallel fiber (PF) synapses involved in producing the erroneous behavior (Albus, 1975; Ito et al., 1982; Marr and Brindley, 1970; Medina and Lisberger, 2008; Yang and Lisberger, 2014). It is proposed that the cerebellum contains internal models which serve feedforward controllers in other brain regions (Wolpert et al., 1998). In response to CF error signals the models would be tuned to modify behavior, through a process of supervised learning. A notable illustration of cerebellar function is observed in adaptation to glass prisms that laterally shift the visual image, leading to initial errors in visually-guided reaching or throwing tasks. Healthy human subjects, as opposed to those with cerebellar damage, typically adjust on the order of 5-10 trials, and retain the adaptation even after the prisms are removed (Baizer et al., 1999; Martin et al., 1996a). However, in this textbook example of cerebellar learning, if the “all-or-nothing” complex spikes (CS), evoked by CF activation, encode error, how do they differentiate between off-target throws in opposite directions and update synapses accordingly to correctly adjust the behavior? Various solutions to this problem have been proposed including modulation of CS amplitude and duration (Najafi et al., 2014a; Rowan et al., 2018; Yang and Lisberger, 2014) or timing (Bouvier et al., 2018; Herzfeld et al., 2018; Ohmae and Medina, 2015) but none of these studies examined population responses to error signals *in vivo* and have not reported CS rate modulation by error magnitude. In this study we find that both error magnitude and sign are reciprocally encoded by the collective CS rate in populations of Purkinje cells.

## Results

### Trial-by-trial adaptation of motor behavior

We trained mice to use a joystick to pull a lick-port close enough to their mouth to consume water droplets (**video 1**). The lick-port’s velocity relative to the joystick (the gain) was adjustable between pulls, enabling the introduction of novel sensorimotor dynamics to which mice had to adapt in subsequent pulls (**Fig. 1A**). After performing around 30 joystick pulls under control conditions (gain = 1), the gain was doubled and mice adapted their behavior accordingly for five consecutive joystick pulls until the gain was returned to control conditions and they recovered from any adaptation for five additional pulls. During recordings, mice completed 603 ± 121 pulls within 60 minutes (mean ± SD; *n* = 13 mice; **Fig. 1B**), enabling the repeated introduction of this sensorimotor perturbation around 50 times per recording session. This pattern allowed detailed monitoring of movement kinematics within an individual recording session including limb position and acceleration (**Fig. 1C-F**) and we monitored how those parameters adapted to and recovered from repeated introduction of the perturbation.

**Figure 1.**
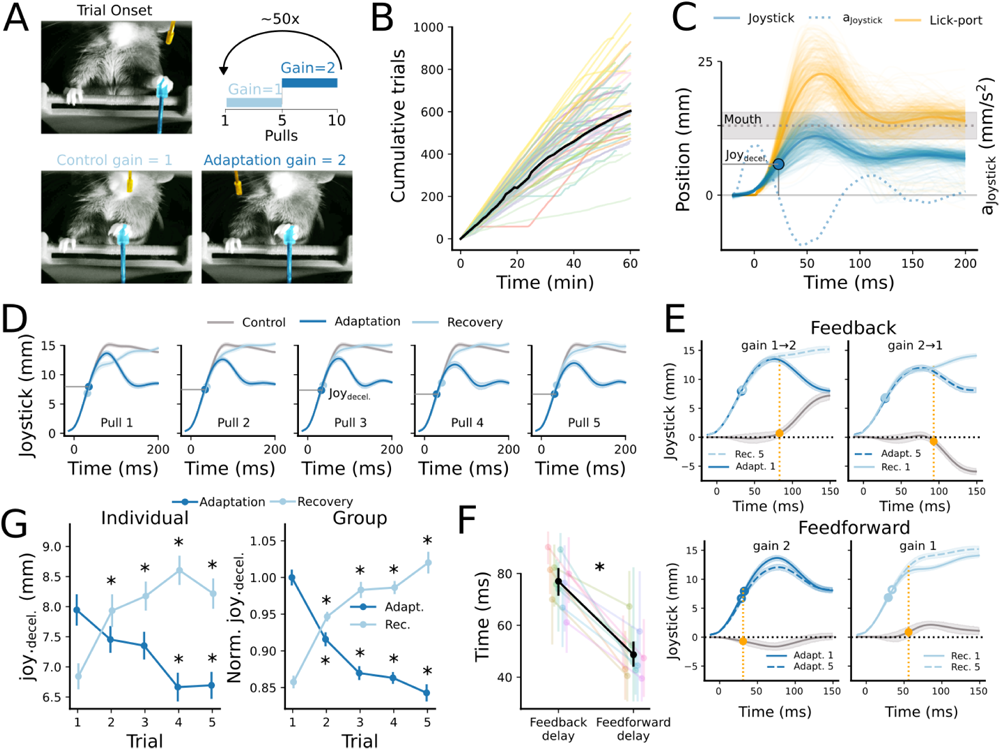
Mice adapt motor commands to quickly respond to imposed sensorimotor perturbations. **A.** Mice were trained to use their left forelimb to manipulate a joystick (blue) that controlled the position of a lick-port (orange) to receive water droplets. A trial structure was imposed whereby mice performed a series of five joystick pulls under control conditions (gain = 1) followed by five pulls with the gain set to 2. This sequence was repeated around 50 times during each recording session, forcing mice to repeatedly adapt (gain 1→2) to and recover (gain 2→1) from the perturbation. **B.** Mice pulled the joystick at a relatively constant rate (10 ± 15 pulls/min) performing 603 ± 121 pulls in 60 minutes (mean ± SD; 64 recordings in 13 mice). **C.** The position of the joystick and lick-port were continuously monitored providing a readout of kinematic parameters such as position (left axis), and acceleration (right axis). The position at which the joystick acceleration was null (joy_decel_; solid blue circle) was tracked and plotted for all pulls in D and G. **D.** Mean trajectories for all pulls during an individual recording session shows the evolution of the joystick kinematics and mean joy_decel_ (blue circles) throughout the series of five pulls in both gain and control conditions (shaded regions, 95% CI). **E.** Upper: The last pull of one condition and the first pull of the next condition begin to deviate from each other 80.4 ms after pull initiation (orange points; feedback delay; *p* < 0.01; Wilcoxon signed-rank test). Grey line indicates the difference between mean trajectories for the first and fifth pulls across conditions (shaded regions, 95% CI). Lower: Same as upper but comparing the 1st and 5th pulls within each gain condition. Orange points are the feedforward delays. **F.** In 105 recording sessions from 13 mice the feedback delay was 80.4 ± 15.0 ms and the feedforward delay was 49.7 ± 18.2 (*p* < 0.01, Wilcoxon signed-rank test; error bars, SEM). **G.** Mean joy_decel_ for all pulls and conditions in the individual recording shown in A-G (left) and for 105 recording sessions from 13 mice (right; *, p < 0.01, Wilcoxon signed-rank test compared to first pull in the condition; error bars, SEM).

To gauge how quickly mice could detect the perturbation and adjust behavior using sensory feedback, we compared average limb trajectories from the last pull of each gain condition to the first pull of the subsequent condition. During these pairs of consecutive pulls, the limb movement started with similar initial trajectories that diverged presumably due to the different sensory feedback in the two conditions. We observed that trajectories began to diverge 80.4 ± 15.0 ms (*n* = 13 mice) after each pull started (**Figure 1E**), giving an indication of the reaction time to sensory feedback. However, the limb trajectories of the first and fifth pulls in the same condition started to diverge much earlier, after just 49.7 ± 18.2 ms (*n* = 13 mice; *p* < 0.01, Wilcoxon signed-rank test comparing feedback and feedforward delays; **Fig. 1E and F**). The adaptation of the initial component of the behavior, well before any influence of sensory feedback, suggests changes to the feed-forward component of the motor command.

To explore the feedforward component of the behavior adaptation in our pulling paradigm we evaluated various kinematic parameters that occurred before the behavior could be influenced by sensory feedback (**fig. S1**). We observed that the limb’s acceleration—a measure proportional to the net force applied to the joystick—ceased 34.4 ± 0.5 ms after pull initiation (example shown in **Fig. 1C-E**), long before any observed effect of sensory feedback. Notably, as illustrated in figures **1D** **and G**, the position where the joystick stopped accelerating (joy_decel._) decreased progressively on a trial-by-trial basis, indicating motor adaptation to the increased lick-port gain (dark blue points in **Fig. 1D and G**). Conversely, upon return to control conditions, this parameter increased incrementally during subsequent pulls as the behavior recovered from the prior adaptation (light blue points in **Fig. 1D and G**). The rapid time course of the adaptation and post-adaptation recovery of this parameter is reminiscent of established cerebellar-dependent behavior paradigms (Martin et al., 1996b; Morton and Bastian, 2006; Smith et al., 2006).

### Sensorimotor mismatch signals in Purkinje cells

The repeated bidirectional adaptation of the behavior within recording sessions allowed us to use two-photon calcium imaging to examine the response of populations of Purkinje cells to the adaptation and recovery phases of the behavior within sessions. Consistent with previous results, imaging 600 x 600 µm sections on the surface of lobules IV/V in mice expressing GCaMP8m targeted to Purkinje cells (using Cre-dependant viral vectors in Jdhu PCP2-cre mice; Witter et al., 2016) revealed long thin regions of interest (ROIs), characteristic of Purkinje cell dendrites, with large spontaneous fluorescence signals, characteristic of CS (1.28 ± 0.21 sp/s, 4577 ROIs from 28 recordings in 7 mice; **Fig 2A and B**; (Gaffield et al., 2019; Kostadinov et al., 2019; Najafi et al., 2014b; Ramirez and Stell, 2016). To begin with, we analyzed whether sensory stimulation associated with the lick-port could affect CS rate in absence of a motor command. We did this by playing back to the mouse the mean position recorded during control pulls, repetitively moving the lick-port in a manner uncoupled from the animal’s actions (**Fig. 2C** **upper panels**). We found that the stimulation caused large modulation of CS rate in individual cells above and below baseline rate (**Fig. 2C and D**) with 119 of 307 cells in a region significantly modulated by the stimulus (*p* < 0.01, Pearson’s χ^2^ test of firing rate compared to baseline in each cell). Surprisingly, this same sensory stimulation—from the whiskers and potentially from olfactory and reward centers—caused significantly less modulation when the mouse was in control of the position (**Fig. 2C-E**; 26/307 cells modulated 33-133 ms after pull initiation; *p* < 0.01, Pearson’s χ^2^ test of firing rate compared to baseline in each cell). This effect, which is similar to that reported for spike rates in the cerebellar nuclei of primates during passive and active head rotation (Brooks and Cullen, 2013), was observed across 25 recordings in 6 animals, with only 16% of cells significantly modulated when the sensory stimulation was controlled by the mouse, whereas 59% were modulated during playback of the lick-port (**Fig. 2E**; *n* = 4194 cells, *p* < 0.01, Pearson’s χ^2^ test of firing rate compared to baseline in each cell). We did not observe any substantial difference between the spontaneous and evoked CS amplitudes or decay times (0.298 [0.295, 0.301] vs 0.305 [0.302, 0.308], ΔF/F0 [95% CI]; 85.4 [84.6, 86.3] vs 84.9 [84.1, 85.8], ms [95% CI]; *n* = 4282 ROIs; **fig. S2**, **tables S1** and **S2**) indicating that CS rate was modulated but there was no effect on the CF-induced Ca^2+^-influx. Overall, this observation indicates that under control conditions, an efferent copy of the motor command actively inhibits the CS evoked by the sensory stimulus.

**Figure 2.**
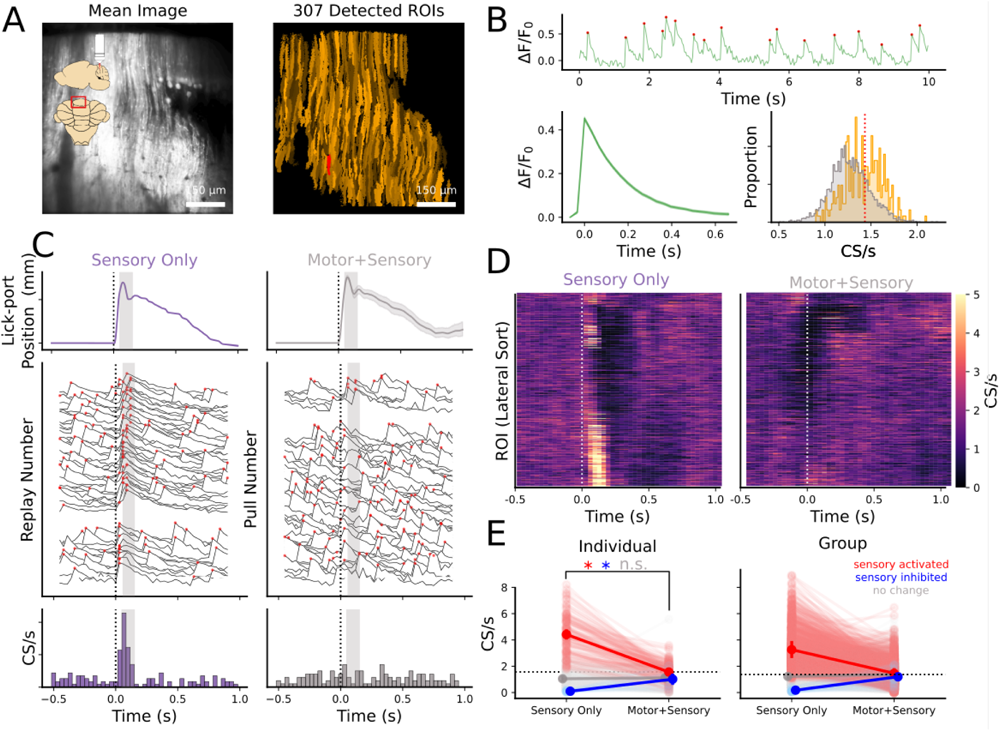
An efferent copy of the motor command cancels sensory evoked CS. **A.** Left: Mean image generated using two-photon imaging of a 600 x 600 um region near the surface of lobules IV/V while a mouse performed the joystick task. Right: 307 ROIs auto-segmented using Suite2P shown in shades of orange. The analysis of the red ROI is presented in subsequent panels. **B.** Upper: 10 seconds of raw fluorescence demonstrates calcium events and their detection (red dots). Lower: mean waveform of all detected events from the same ROI and distribution of event rate for all ROIs (gray), the ROIs of the region recorded in A (orange), and the red ROI from A (red dotted line). **C.** Upper: Mean lick-port position played back for the sensory only condition (violet) and lick-port trajectory recorded when the mouse moved the joystick (gray). Middle: Fluorescence and detected events (red dots) for individual playback of the movement (left) and individual pulls (right) from the same ROI, grouped by event presence during a 100 ms window during the movement (shaded region). Lower: PSTHs show modulation when the lick-port is played back but not when the mouse is in control of the movement. **D.** CS rates of all 307 ROIs shown in A. **E.** Comparison of CS rates for all ROIs that were increased (red) or decreased (blue) in the sensory only condition and the response of those same cells when the mouse controlled the lick-port (motor + sensory). Individual recording session shown in A (left) as well as for all mice (right; *, *p* < 0.01, Wilcoxon signed-rank test; *n* = 25 recordings in 6 mice).

If CS evoked by lick-port movement are inhibited by an efferent copy of the motor command, the command would have to be predictive of its sensory consequences. As shown in figure **1E**, on the first pull in one condition—mice unaware that the gain was changed between pulls—begin to execute the same movement as in the previous condition. Consequently, on the first pull in either condition there is a mismatch between the expected and actual sensory reafference which progressively realigns as the mouse adapts its behavior to the new lick-port gain. To determine if this mismatch was signaled to the cerebellum, we focused on the activity evoked during the movement and analyzed CS rates in a 100 ms window after pull initiation. In an individual recording, we observed dendrites with CS rates that were significantly modulated (91/159 cells; *p* < 0.01, Pearson’s χ^2^ test of firing rate compared to baseline in each cell; **Fig. 3D**) during the first pulls after the gain was increased and that modulation trended back towards the baseline firing rate as the mouse adapted its motor command to the new sensorimotor dynamics in subsequent pulls (*p* < 0.01, Wilcoxon signed-rank test compared to the first pull; **Fig. 3D**). Surprisingly, when the gain of the lick-port movement was then switched back to control conditions and the mouse attempted to use the motor command that was now adapted to the perturbed condition, an inhibition of the CS rate was revealed in these same cells, that also trended back towards baseline in subsequent pulls (*p* < 0.01, Wilcoxon signed-rank test compared to the first pull; **Fig. 3D**). As shown in figure **3B** this effect was observed in many cells throughout the recording region. When we analyzed all cells across 17 recordings in 7 mice, that were significantly modulated by the first pull in the adaptation condition (**Fig. 3D**), we found that this modulation decreased across subsequent pulls (-0.53 CS/s; *p* < 0.01, Wilcoxon signed-rank test comparing the rates evoked during the first and fifth pulls). Furthermore, cells that were activated when the mouse was adapting to the perturbation, were inhibited when the gain was restored to control conditions (+0.26 CS/s; *p* < 0.01, Wilcoxon signed-rank test; **Fig. 3D**). Together, these observations suggest that a mismatch between the expected and perceived reafference causes bidirectional modulation of CS firing rate above and below the baseline rate, and this modulation reflects the bidirectional adaptation of behavior.

**Figure 3.**
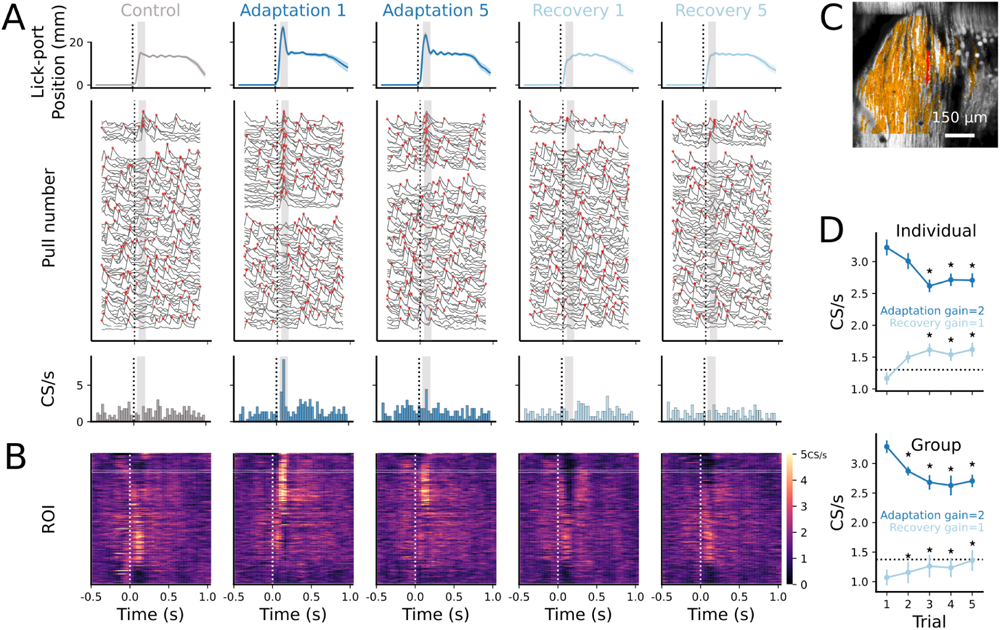
Reciprocal modulation of CS rate by behavioral adaptation. **A.** Upper: mean lick-port trajectories during one recording session for the control, as well as the first and fifth pulls of the joystick in each condition. Middle: Fluorescence and detected events (red dots) for the red highlighted cell in C. Lower: PSTHs show an increased rate on the first pull in the gain = 2 condition that decreases by the fifth pull, and an inhibition below baseline rate on the first pull of the gain = 1 condition that trends back to baseline by the fifth pull. Shaded regions indicate the 100 ms window during the movement used in D. **B.** CS rates of all ROIs shown in C organized laterally show groups of ROIs that are activated by the switch to the gain = 2 condition and inhibited by the switch to the gain = 1 condition and vice versa. White box highlights the cell shown in A and C. **C.** Mean image with 159 ROIs (shades of orange) and the highlighted cell in red. **D.** CS rates across five pulls for all ROIs that were significantly activated by the first pull in the gain = 2 condition (dark blue; *,*p* < 0.01, Pearson’s χ^2^ test compared to the first pull). Those same ROIs in the gain = 1 condition were inhibited (light blue). This was the case in the individual recording shown in E (upper) as well as across 17 recordings in 7 mice (lower; error bars, SEM).

### CS rate in parasagittal bands is modulated by sign and magnitude of sensorimotor mismatch

To determine the extent and reproducibility of the CS rate modulation we imaged multiple regions within and across seven different mice. In figure **4A** we show data from 4 recordings of the same mouse performed on separate days with each recording outlined in gray. The mean CS rate is plotted for all dendrites during a 100 ms window after the start of the first pull in each condition. This representation demonstrates cells with similar mean responses organized in parasagittal bands across the entire recording area and individual bands with opposing responses to each gain condition spanning across recording sessions/regions. To further investigate the spatial organization of responding Purkinje cells, we used *k*-means clustering of the mean CS response of each cell during pulls in order to identify cells that responded similarly (materials and methods). Clustering on the activity alone also identified groups of cells organized in parasagittal bands. This robust pattern was observed in all mice recorded (4 examples shown in **Fig. 4B**). Although we restricted clustering to activity occurring during a one second window following pull initiation, the mean CS rates for clusters often displayed large modulation prior to pull initiation (**Fig. 4C**). This observation is consistent with previous reports (Calame et al., 2023; Wagner et al., 2021) but as we are interested in error signals evoked by reafference, we focused on the modulation after pull initiation. When plotting the mean PSTH from four mice, for select parasagittal bands displaying modulation of CS rate immediately following pull initiation (shaded regions in **Fig. 4C**), we observe maximum modulation—excitatory or inhibitory depending on the cluster—on the first pull in a condition that trended towards the baseline rate as the behavior adapted to the new sensorimotor dynamics (**Fig. 4C**). Furthermore, these plots demonstrate that unlike activity modulated prior to movement initiation, the CS rate observed when the lick-port was moving during the 100 ms following pull initiation was reciprocally modulated by the gain condition.

**Figure 4.**
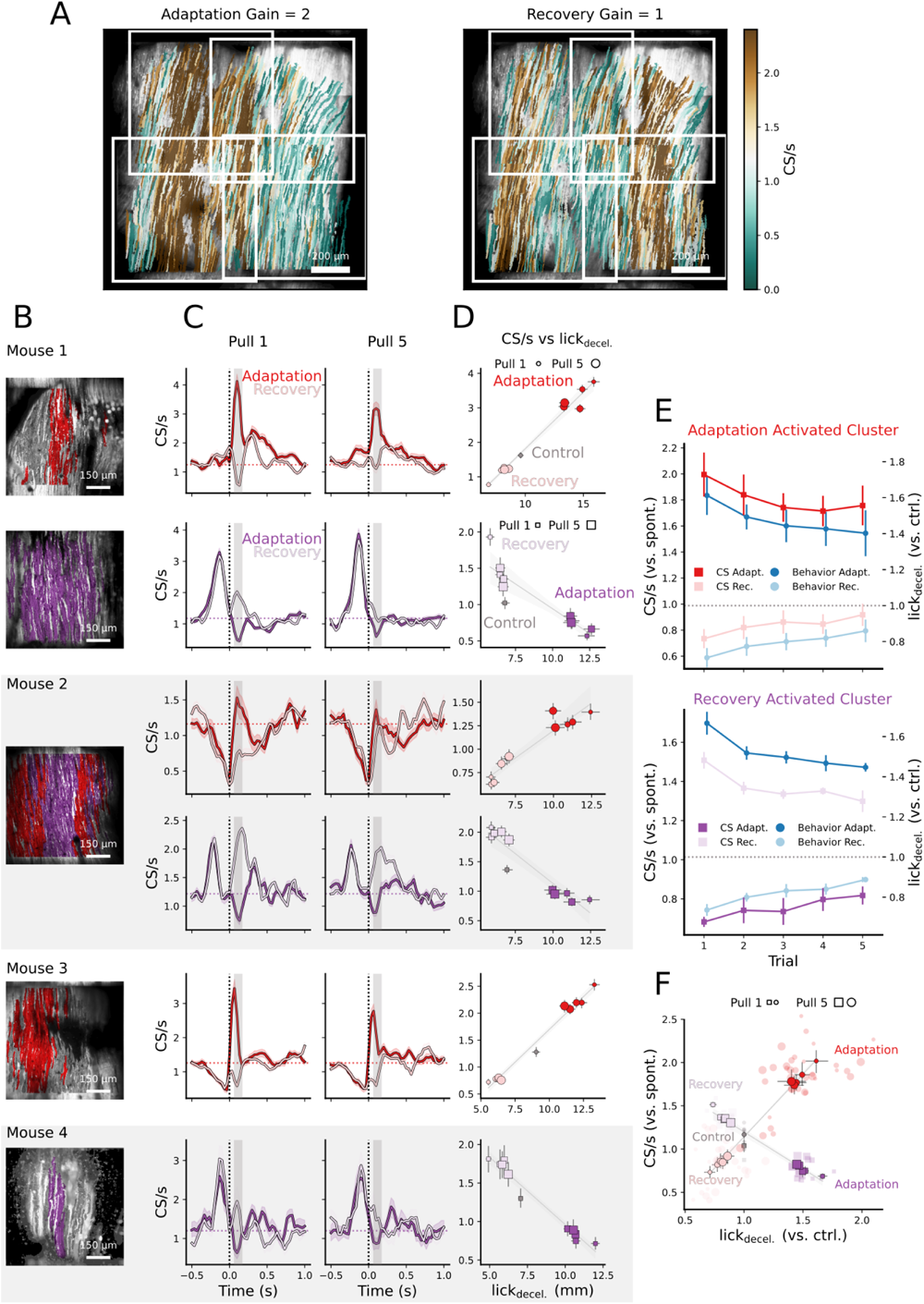
Parasagittally organized bands of cells are reciprocally modulated by error amplitude. **A.** Mean CS rates in a 100 ms window after pull initiation during first pulls in the adaptation (left) and recovery conditions (right) across four neighboring regions (outlined in white) of lobule IV/V of one mouse. There exist bands of activity that are reciprocally modulated in each condition. **B.** Mean images from four of the seven different mice. *k*-means clustering of mean activity reveals parasagittal clusters of ROIs that increase (red) or decrease (violet) CS rate in response to the sensorimotor perturbation. **C.** Rolling mean (window size = 3) of PSTHs of mean CS rates for all ROIs of the clusters indicated in B demonstrate that clusters which are positively modulated by the perturbation (red), are inhibited when the gain returns to control conditions (pink). The clusters that display decreased CS rates in response to the perturbation (violet), are excited upon return to control gain (light violet). PSTHs for the fifth pull are less modulated than the first. Shaded boxes are the 100 ms windows used in D-F. **D.** Mean CS rate vs. the deceleration point of the lick-port (lick_decel_) for every pull grouped by pull number (marker size) and condition (saturation). This shows a clear correlation within behavior sessions between lick-port position and CS rate. **E.** Upper: Group data of CS rates normalized to baseline (left axis) for all clusters positively modulated during adaptation pulls and normalized lick_decel_ (right axis), dark colors depict CS rates during adaptation pulls, and lighter colors depict recovery pulls. This shows the reciprocal rate and behavior modulation across conditions and the evolution towards control across subsequent pulls (17 sessions in 7 mice). Lower: Same as the upper panel but for clusters that were inhibited during adaptation pulls (11 sessions in 3 mice). **F.** Same as D for all recording sessions normalized to the baseline rate and control lick_decel_ of the recording. Red and violet markers indicate excited and inhibited clusters, during adaptation pulls, respectively. The first pulls (smallest markers) in either condition have the biggest effect with subsequent pulls trending towards the control positions/rates (error bars, SEM).

The above results are consistent with a mechanism by which the sign and magnitude of CS modulation encode sensorimotor error evoked by a difference between the expected and perceived reafference associated with a motor command. Since the reafference is a function of the lick-port trajectory, we quantified it using the deceleration point of the lick-port (lick_decel_)–similarly to joy_decel_ (**Fig. 1**)–and any error would be the difference from unperturbed control pulls. We plotted the responses of modulated clusters against lick_decel_, grouped by condition and pull number, and demonstrate that the sign of the modulation, compared to baseline firing, is determined by the condition, and thus the sign of lick_decel_. Furthermore, within each condition the magnitude of CS modulation scales with the magnitude of lick_decel_ (**Fig. 4D**).

The relationship between error amplitude and CS rate is consistent in the seven mice recorded for bands that were activated (**Fig. 4E, upper**) as well as those that were inhibited (**Fig 4E, lower**) when the lick-port overshoots the mouth upon the switch from control to gain = 2 conditions. Plotting the activated and inhibited clusters for all mice together in **Fig. 4F** highlights the consistency of the reciprocal relationship between CS rate and the sign of the error, with CS rate augmented by the error evoked in one gain condition and inhibited by the opposite error created upon transition to the other condition. The relationship between error magnitude and CS rate modulation is also evident here, the first pulls in each condition show the largest deviation from control pulls evoking the largest modulation of CS rate, while subsequent pulls trend towards baseline rates as the behavior trends towards the control behavior.

## Discussion

These results demonstrate that together, CS within parasagittal bands of Purkinje cells encode direction and amplitude of a sensorimotor error. Bands that increase CS rates above baseline when mice initially pull the joystick too far, are inhibited below baseline when the joystick is not pulled far enough, and vice versa. Synapses onto Purkinje cells have been extensively studied and shown to undergo opposing signs of plasticity depending on their timing with respect to a CS (Bouvier et al., 2016; Coesmans et al., 2004; Ito et al., 1982; Lev-Ram et al., 2002; Safo and Regehr, 2008; Sarkisov and Wang, 2008). Together with the results above, this provides a potential mechanism by which inputs onto bands of Purkinje cells involved in this behavior could be collectively potentiated or inhibited, with the direction of this plasticity depending on the result of the behavior and whether the band ultimately affected the extensor or flexor component of the behavior.

Similarly to the trial-by-trial adaptation of human patients attempting to throw with perturbed vision, mice in our joystick paradigm display behavior modifications after only one pull in response to novel sensorimotor dynamics. This is in contrast to typical LTP/LTD induction protocols referenced above which require hundreds of CF/PF pairings to observe synaptic changes. However, in slice physiology experiments, synaptic changes need to be large enough to record in individual cells, whereas the smallest change induced by a single pairing could have a significant downstream effect when induced collectively in the subpopulation of cells we observe here.

Although our study does not directly reveal the mechanism by which CF/CS modulation occurs, we show that CS evoked by sensory stimulation are reduced when that stimulation is self-generated. Further experiments are required to determine whether the cancellation occurs at an early stage of sensory processing (Singla et al., 2017) or involves functional units of the cerebellum (Oscarsson, 1979).

## Acknowledgments

We are grateful to the following for discussions throughout the project and/or comments on the manuscript: Philippe Ascher, Céline Auger, Boris Barbour, Isabel Llano, and Alain Marty.

## Funding

Agence nationale de la recherche N° ANR-19-CE37-0011-01

## Author contributions

Conceptualization: BMS

Methodology: BMS

Investigation: VN, BMS

Visualization: VN, BMS

Funding acquisition: BMS

Project administration: BMS

Supervision: BMS

Writing – original draft: VN, BMS

Writing – review & editing: VN, BMS

## Competing interests

Authors declare that they have no competing interests.

## Supplementary Materials

### Materials and Methods

#### Animal Care & Housing

All procedures were approved by the Université Paris-Cité Animal Ethics Committee and were in accordance with the European Community Council Directive of 22 September 2010 (2010/63/UE) and the French National Committee (87/848) for the care and use of laboratory animals. All efforts were made to minimize animal suffering and to reduce the number of animals used. All animals were housed in the animal facility of the Université Paris-Cité (Paris, France) in a temperature-controlled room (22 ± 1°C) with a 12 h light/dark cycle (lights on at 7:00 a.m.) and had *ad libitum* access to food. We used mice expressing Cre-recombinase specifically in PC under the control of the L7 promoter (B6.Cg-Tg(Pcp2-cre)3555Jdhu/J; The Jackson Laboratory, Bar Harbor, ME, USA).

Animals were housed in groups of 2-5 animals per cage. The cages were enriched with a cardboard tube and a plastic house. Animals were chronically water-deprived for up to 5 days at a time during experimental periods. During water deprivation, animals fulfilled the majority of their hydration requirements during the behavioral task. Animals were weighed daily, after each experiment, and their weight was maintained at 85-90% of their free-ingestion weight with supplemental water. The animals were monitored daily for signs of distress and were euthanized if they showed signs of distress or if they lost more than 20% of their free-ingestion weight.

#### Surgical Procedures

Animals used in all experiments were implanted with a stainless steel headpiece for head-restraint. Animals which were used for calcium imaging additionally underwent stereotaxic, intracranial viral injection, craniotomy and cranial window implantation. All procedures were performed consecutively on the same day.

#### Preparation & Anaesthesia

Anaesthesia was first induced by placing the animal in a chamber with 3% v/v isoflurane in oxygen. Once the animal was anesthetized, it was placed in a stereotaxic frame (Kopf Instruments, Tujunga, CA, USA) using ear bars and a tooth bar. The anesthesia, now being delivered via a mask, was reduced to 1.5% v/v isoflurane. Anesthesia was regulated via a Univentor 410 anesthesia unit (Univentor Ltd., Zejtun, Malta). The animal sat atop a heating pad and the temperature was monitored with a rectal probe. The animal’s eyes were protected with a thin layer of eye ointment (Optigel; Covalis, Tourville-la-Rivière, France). The animal was given a subcutaneous injection of 0.1 mg/kg buprenorphine (Buprécare; Axience SAS, Pantin, France). The surgical zone was prepared by shaving followed by disinfection with 70% v/v ethanol and povidone-iodine solution (Vétédine; Vetoquinol, Lure, France) then by a subcutaneous injection of 100 µL lidocaine (Xylovet; Centravet, Dinard, France).

#### Headpiece Implantation

A circular section (diameter = 10 mm) of skin was removed over the occipital bone and the skull was cleaned by scraping with a no. 24 scalpel blade followed by application of 3% v/v hydrogen peroxide which was left for 30 seconds and then removed. Machined stainless steel headpieces were of our own design. The headpieces were attached to the skull using dental cement (Super-Bond C&B, Sun Medical, Shiga, Japan).

#### Craniotomy & Viral Injection

Animals used for calcium imaging experiments expressed the calcium indicator GCaMP8m (Zhang et al., 2023) in PC in a specific manner under the control of the Pcp2/L7 promoter. In order to achieve this, the L7-cre-2 mice were injected with a Cre-dependent viral vector (#162378; Addgene, Watertown, MA, USA), concurrently with a vector to express tdTomato (#51503; Addgene, Watertown, MA, USA), also in a Cre-dependent manner, which served as a reference for non-calcium related changes in fluorescence. Firstly, a 3 x 3 mm square craniotomy over the occipital bone was performed using a dental drill with a 0.3 mm dental burr (BUSCH & CO. GmbH & Co. KG, Engelskirchen, Germany). A solution containing the two viral vectors was prepared in phosphate-buffered saline (PBS) to concentrations of 6.5 x 10^11^ vg/mL and 2.5 x 10^12^ vg/mL for the GCaMP8m and tdTomato vectors respectively. The solution was loaded into a 25 µL Hamilton syringe (Hamilton Company, Reno, NV, USA) mounted in a stereotaxic syringe pump (Legato 130; KD Scientific, Holliston, MA, USA). The viral vectors were injected at a rate of 100 nL/min using a beveled glass pipette, at a depth of 300 µm below the cortical surface in lobules IV/V of the cerebellar vermis. The pipette was left in place for 10 minutes after the injection to allow for diffusion of the viral vector.

#### Cranial Window

A glass cranial window was implanted over the craniotomy to seal the brain from the environment yet allow for optical access. The cranial window was composed of multiple layers of glass glued together with UV-curing optical adhesive (NOA61, Norland Products Inc., Cranbury, NJ, USA). A single upper layer of 3.2 x 3.2 mm glass was glued to 3 lower layers of 3 x 3 mm glass, each layer was hand-cut from 200 µm thick glass coverslips (VWR International, Radnor, PA, USA). The lower layers of glass were inserted into the craniotomy, pressing on the brain and the upper layer was glued to the skull with dental cement (Super-Bond C&B; Sun Medical, Shiga, Japan).

#### Experimental Apparatus & Behavioural Task

Water-restricted, head-restrained animals manipulated a lick-port by controlling the position of a joystick. The task required positioning of the lick-port in front of the animal’s mouth to consume droplets of water. The animal was head-restrained, and a tube was placed on top of it to restrain movement of the body and hindlimbs while leaving the forelimbs free to operate the joystick. The joystick was a 13 cm long 3D printed plastic rod which was attached perpendicularly atop an 8 mm steel axle which was supported by two cartridge bearings. One end of the axle was mated to an AMT112S rotary encoder (CUI Inc., Tualatin, OR, USA) which we used to track the position of the joystick. The joystick’s movement was limited to an arc of 25 mm. An elastic band pulled the joystick to the home position, the left limit of the arc.

The lick-port position was set by a NEMA 17 pattern stepper motor (QSH4218; Trinamic Motion Control GmbH & Co. KG, Hamburg, Germany). The lick-port was attached to a profiled band and was moved laterally by the turning of a pulley wheel, driven by the stepper motor. The stepper motor was controlled by a Watterott TMC2100 stepper motor driver (Watterott Electronic GmbH, Germany) which in turn was controlled by a Teensy 4.1 microcontroller (PJRC, LLC, Sherwood, OR, USA). The Teensy was programmed to receive TTL pulses from the rotary encoder and to convert these pulses into stepper motor commands. The Teensy was programmed to move the lick-port in response to joystick movement in a hybrid open/closed-loop manner. The gain of the lick-port could be modulated via the program on the Teensy.

#### Behavioral Habituation

Prior to initial training, animals underwent 24 hr of water deprivation. The animal was initially presented with an easier version of the task:. The home position was initially set 5 mm away from the animal’s midline and as the animal began performing the task, the experimenter would progressively a) retract the home position away from the animal and b) decrease the reward size. Training required approximately 1 week and animals were considered experts once they were able to satisfy their daily hydration requirements with the home position set at 12 mm from centreline (approximately 600 pulls, 1 µL dispensed per pull).

#### Adaptation Task

Once the animals were trained to expert level they were executing a well refined motor program which resulted in stereotyped trajectories of the joystick and lick-port. The animals were then subjected to the adaptation task, wherein the gain of the lick-port was constantly switched between 1 and 2, which required them to constantly adapt their motor program. The experiment was divided into adaptation (gain = 2) and recovery (gain = 1) blocks of 5 trials each. Mice only performed one recording session per day.

#### Playback Condition

As the animal moves the joystick, the joystick in turn causes the lick-port to displace. We introduced the playback condition to produce the same movement as that caused by the animal’s own movements, but in an uncoupled manner. During the block of control trials, each trajectory was recorded and the mean trajectory was stored as a vector on the Teensy microcontroller. During a separate playback block, the lick-port was uncoupled from the joystick and instead the stored vector was used to control its position. The lick-port would “playback” at random intervals between 2 s and 10 s.

#### Calcium Imaging

With animals expressing GCaMP8m fluorescent calcium indicator, and tdTomato fluorescent protein, we recorded the activity of PC in the cerebellar vermis using two-photon laser scanning microscopy (2PLSM). Both fluorophores were excited by pulsed 910 mm wavelength light, emitted from a Spectra Physics Mai Tai Ti:Sapphire laser (< 100 fs pulse width; 80 MHz repetition rate; MKS Instruments Inc., Andover, MA, USA). Laser was focused through an Olympus XLUMPLFLN20XW water-immersion objective (20 x; NA = 1; Olympus Corporation, Tokyo, Japan), and average power during recordings, measured at the objective, was approximately 100 mW. Emitted fluorescence was captured though the same excitation pathway, and separated via a dichroic mirror. The emitted green and red photons were then separated with another dichroic mirror along with appropriate filters and each green and red channel was recorded separated via independent photomultiplier tubes (PMT2101/M; Thorlabs Inc., Newton, NJ, USA). Scanning of the laser beam over a 600 x 600 µm surface was accomplished using a Sutter Resonant Scan Box (RESSCAN; Sutter Instrument Company, Novato, CA, USA). Frames were acquired at 30 Hz, scanning 512 lines per frame.

#### Data Acquisition

Acquisition of signals from the PMTs, scan control, as well as acquisition of behavioral data, were performed using a pair of NIDAQ systems (National Instruments Corporation, Austin, TX, USA). Imaging was handled by ScanImage software (Vidrio Technologies LLC. Ashburn, VA, USA) through a NI PCIe-8361/PXIe-1073: scan control signals were generated in ScanImage and sent to the RESSCAN via a NI PXIe-6341, PMT signals were amplified before being digitized at 30 kHz via a National Instruments PXIe-7961 FlexRIO FPGA (programmed by ScanImage), and then sent to be processed by ScanImage via a second NI PXIe-6341. ScanImage saved the data stream as TIFF files. The behavioral data (digital) was sampled at a rate of 100 KHz via a NI PCIe-6353 using WaveSurfer (Janelia Research Campus, Ashburn, VA, USA), which stored the binary data stream in a HDF5 container. The two data streams were synchronized via a TTL pulse sent from the Teensy microcontroller to both NIDAQs at the start of each trial, and then later aligned by the unique time delta between trials.

#### Data Analysis

All data were analyzed with custom scripts written in Python. The number of clusters (*k*) for *k*-means clustering was determined manually and then later validated using the silhouette method (Rousseeuw, 1987). CS amplitude was calculated as the F value of the cell in the frame where the spike was detected, subtracted by its value in the preceding frame (F_0_).

#### Detection & Curation of ROIs

ROIs corresponding to PC dendrites were detected using Suite2P (Pachitariu et al., 2017), which was configured to use activity based detection. To classify the ROIs we built a regression model using the least absolute shrinkage and selection operator (LASSO; Santosa and Symes, 1986; Tibshirani, 1996). The model was trained on 624 ROIs, which we manually labeled as either “good” or “bad”, using features of ROI shape from Suite2P (size, aspect ratio etc.), transformations engineered from the ROI’s fluorescence signal (high/low-pass filtering), as well as various descriptive statistics (mean, SD, CV, skew, kurtosis).

#### Spike Detection

Spikes were detected by thresholding the deconvolved fluorescence traces of each ROI. Deconvolution was performed using the built-in class of the Suite2P package, which itself is based on the OASIS algorithm (Friedrich et al., 2017). The thresholding was performed after division of the deconvolved traces by median fluorescence, to account for differences in baseline fluorescence, and then by their 99^th^ percentiles in order to account for differences in signal-to-noise ratio.

---

**Figure S1.**
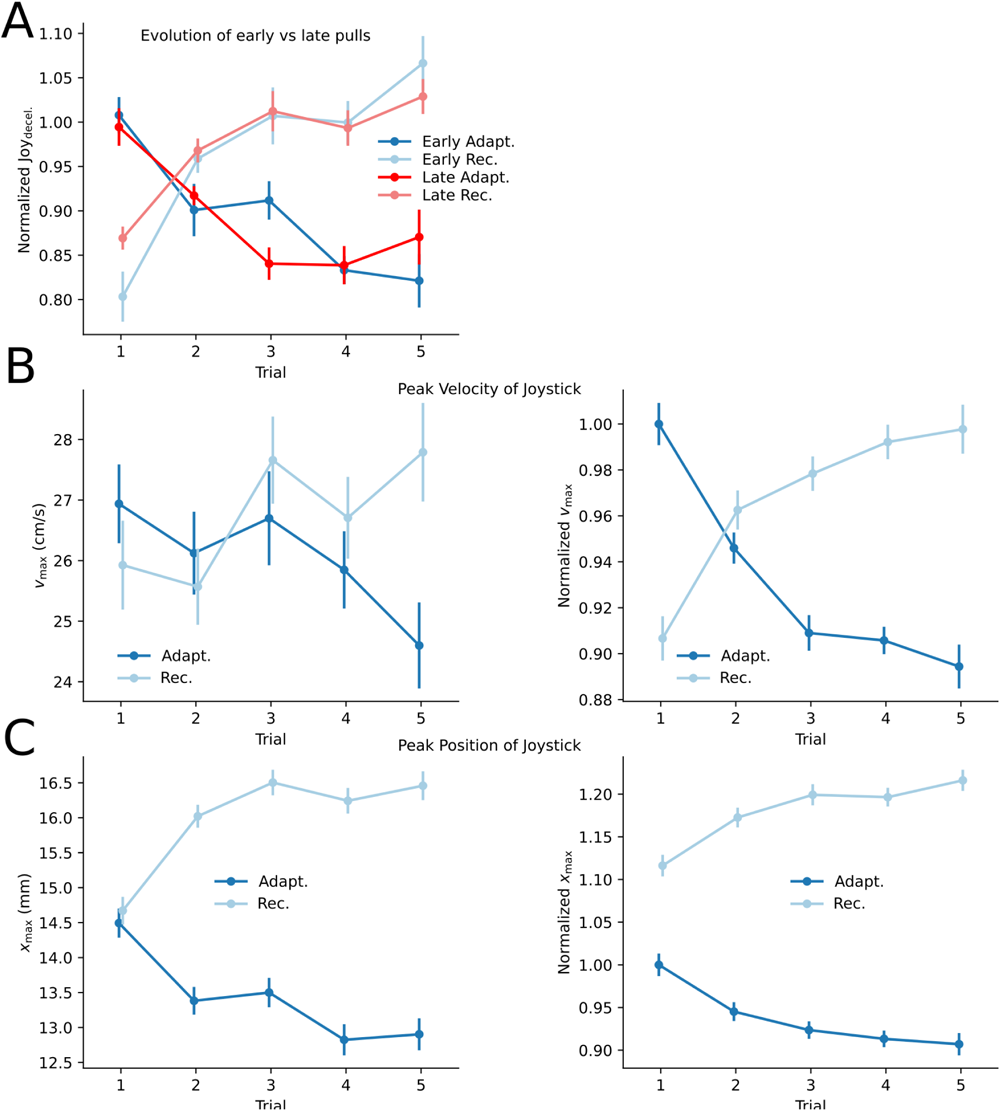
Analysis of various behavior parameters. **A.** Joy_decel_ normalized to the first pull for group data (105 recordings across 13 mice). The similar evolution across the first 250 pulls (blue) and the next 250 pulls (red) within a session shows that learning is the same at the beginning and end of a recording session. **B.** Maximum velocity of the limb/joystick for the example recording in Figure 1 (left) and for all animals (right). **C.** The maximum position of the joystick in the 100 ms following pull initiation for the example recording in Figure 1 (left) and for all animals (right).

**Figure S2.**
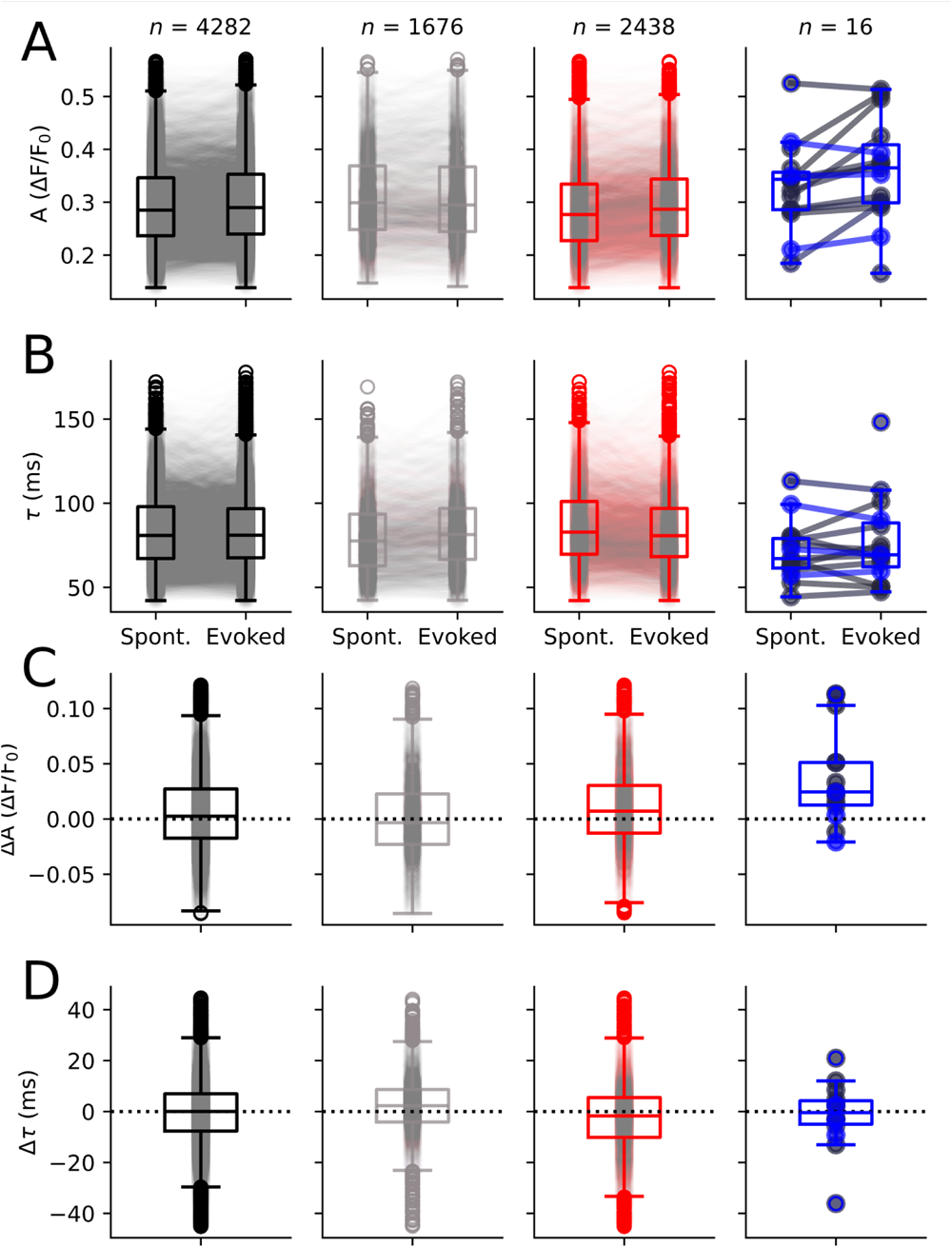
Kinetics and amplitudes of spontaneous CS versus those that were evoked by the sensory alone condition. **A.** Individual recordings (circles) and box-plot distributions of amplitude (A) of spontaneous and evoked CS across all cells (black), those that increased (red), decreased (blue) or showed no change (gray) in firing rate in response to the lick-port movement. **B.** Same as A for the CS decay times (τ). **C.** The difference in amplitude between evoked and spontaneous CS for the same conditions as above. **D.** Same as C for the decay times. Dashed lines indicate no change.

**Table S1.** Kinetics and amplitudes of spontaneous CS versus those that were evoked by the sensory alone condition. A, amplitude; τ, decay time constant.

|  | All |  | Unmodulated |  | Excited |  | Inhibited |  |
| --- | --- | --- | --- | --- | --- | --- | --- | --- |
|  | Spont. | Evoked | Spont. | Evoked | Spont. | Evoked | Spont. | Evoked |
| **A ($\Delta F/F_0$)** | 0.298 | 0.304 | 0.313 | 0.313 | 0.288 | 0.299 | 0.317 | 0.375 |
| **A 95%CI** | (0.295, 0.301) | (0.302, 0.308) | (0.308, 0.318) | (0.308, 0.318) | (0.284, 0.291) | (0.295, 0.302) | (0.266, 0.369) | (0.315, 0.435) |
| **$\tau$ (ms)** | 85.4 | 84.9 | 82.2 | 85.6 | 87.728 | 84.6 | 78.6 | 75.1 |
| **$\tau$ 95% CI** | (84.6, 86.3) | (84.1, 85.8) | (80.8, 83.6) | (84.1, 87.1) | (86.7, 88.7) | (83.5, 85.5) | (61.0, 96.1) | (60.7, 89.5) |

**Table S2.** Differences between spontaneous and evoked CS from Table S3. A, amplitude; τ, decay time constant.

|  | All | Unmodulated | Excited | Inhibited |
| --- | --- | --- | --- | --- |
| **$\Delta A$ ($\Delta F/F_0$)** | 0.006 | -0.000 | 0.011 | 0.058 |
| **$\Delta A$ 95% CI** | (0.005, 0.008) | (-0.001, 0.001) | (0.010, 0.012) | (0.055, 0.060) |
| **$\Delta \tau$ (ms)** | -0.504 | 3.409 | -3.209 | -3.461 |
| **$\Delta \tau$ 95% CI** | (-1.022, 0.014) | (2.561, 4.258) | (-3.862, -2.556) | (-10.71, 3.788) |

## Notes

### Competing Interest Statement

The authors have declared no competing interest.

